# The irradiated brain microenvironment supports glioma stemness and survival via astrocyte-derived Transglutaminase 2

**DOI:** 10.1101/2020.02.12.945329

**Authors:** Tracy J. Berg, Carolina Marques, Vasiliki Pantazopoulou, Elinn Johansson, Kristoffer von Stedingk, David Lindgren, Elin J. Pietras, Tobias Bergström, Fredrik J. Swartling, Valeria Governa, Johan Bengzon, Mattias Belting, Håkan Axelson, Massimo Squatrito, Alexander Pietras

## Abstract

The tumor microenvironment plays an essential role in supporting glioma stemness and radioresistance, and following radiotherapy, recurrent gliomas form in an irradiated microenvironment. Here, we found that astrocytes, when pre-irradiated, increased stemness and survival of co-cultured glioma cells. Tumor-naïve brains increased reactive astrocytes in response to radiation, and mice subjected to radiation prior to implantation of glioma cells developed more aggressive tumors. We identified extracellular matrix derived from irradiated astrocytes as a major driver of this phenotype, and astrocyte-derived transglutaminase 2 (TGM2) as a promoter of glioma stemness and radioresistance. TGM2 levels were increased after radiation in vivo and in recurrent human glioma, and TGM2 inhibitors abrogated glioma stemness and survival. These data suggest that irradiation of the brain results in the formation of a tumor-supportive microenvironment. Therapeutic targeting of radiation-induced, astrocyte-derived extracellular matrix proteins may enhance the efficacy of standard of care radiotherapy by reducing stemness in glioma.

## Introduction

Glioblastoma multiforme (GBM) is the most aggressive and most common (highest grade) glioma, with less than a 10% five year survival rate (Stupp et al. 2009). GBM is typically treated by surgical resection, radiation, and chemotherapy, yet nearly all tumors recur after treatment (Huse and Holland 2010). Recurrent tumors generally form within or overlapping the original tumor volume (Hess et al. 1994) and within a gliotic region corresponding to the initial field receiving radiation, suggesting that the recurring tumor forms within the irradiated brain microenvironment. The recurred tumors have limited treatment options, and the mechanisms underlying therapeutic resistance are incompletely understood.

Recurrence of GBM is often attributed to radioresistant glioma cells. The radioresistant tumor cell phenotype appears to be related to tumor stemness, in that tumor cells surviving radiotherapy display characteristics of stem cells, such as enhanced self-renewal and increased drug efflux capabilities, as compared to radiation-sensitive tumor cells.(Bao et al. 2006; Lathia et al. 2015) Understanding which factors promote stemness and treatment resistance of glioma cells is of intense interest in developing new therapies in glioma. Microenvironmental cues such as hypoxia (Li et al. 2009), extracellular matrix proteins (Lathia et al. 2012; Motegi et al. 2014; Pietras et al. 2014; Farace et al. 2015; Pointer et al. 2017; Barnes et al. 2018; Li et al. 2018; Yu et al. 2018), or growth factors secreted by stromal cells (Wang et al. 2018; Hide et al. 2019) may be sufficient to induce tumor cell stemness and therapeutic resistance. Microenvironmental regulation of tumor stemness is further supported by the finding that stem-like tumor cells are enriched in specific GBM tumor niches, primarily the perivascular niche (PVN) and hypoxic compartments (Calabrese et al. 2007; Li et al. 2009; Hambardzumyan and Bergers 2015). These niches can contain a variety of stromal cell types including astrocytes, microglia, endothelial cells, and pericytes (Quail and Joyce 2017). While several of these cell types contribute to glioma aggressiveness (Pyonteck et al. 2013; Mega et al. 2020), little is known about how these cells respond to radiotherapy, and how they in turn might affect the associated glioma cells after irradiation. We sought to determine whether irradiation of the brain tumor microenvironment might affect the phenotype and therapeutic response of glioma cells. We identified astrocyte-derived transglutaminase 2 (TGM2) as a potential radiation-induced modifier of the tumor microenvironment, which protected against radiation-induced glioma cell death, and may thus serve as a potential therapeutic target in GBM.

## Results

### Irradiated astrocytes modify the extracellular matrix to promote glioma cell stemness and radiation resistance

We devised a co-culture scheme with which we could rapidly test a variety of control or irradiated stromal cells for their ability to increase one of the indicators of stemness in glioma cells, the side population (SP).(Bleau et al. 2009) To do this, we pre-irradiated four stromal cell types before co-culturing with non-irradiated PDGFB-induced glioma primary cells (PIGPC) derived from RCAS-*PDGFB*-induced gliomas in *Nestin-tv-a Ink4a/Arf*^*-/-*^ mice. Non-irradiated PIGPC were added to the pre-irradiated cells for co-culture for 48 h, followed by an SP assay. The screen design is outlined in Figure 1A (Fig S1A-B for gating strategy).

**Figure 1.**
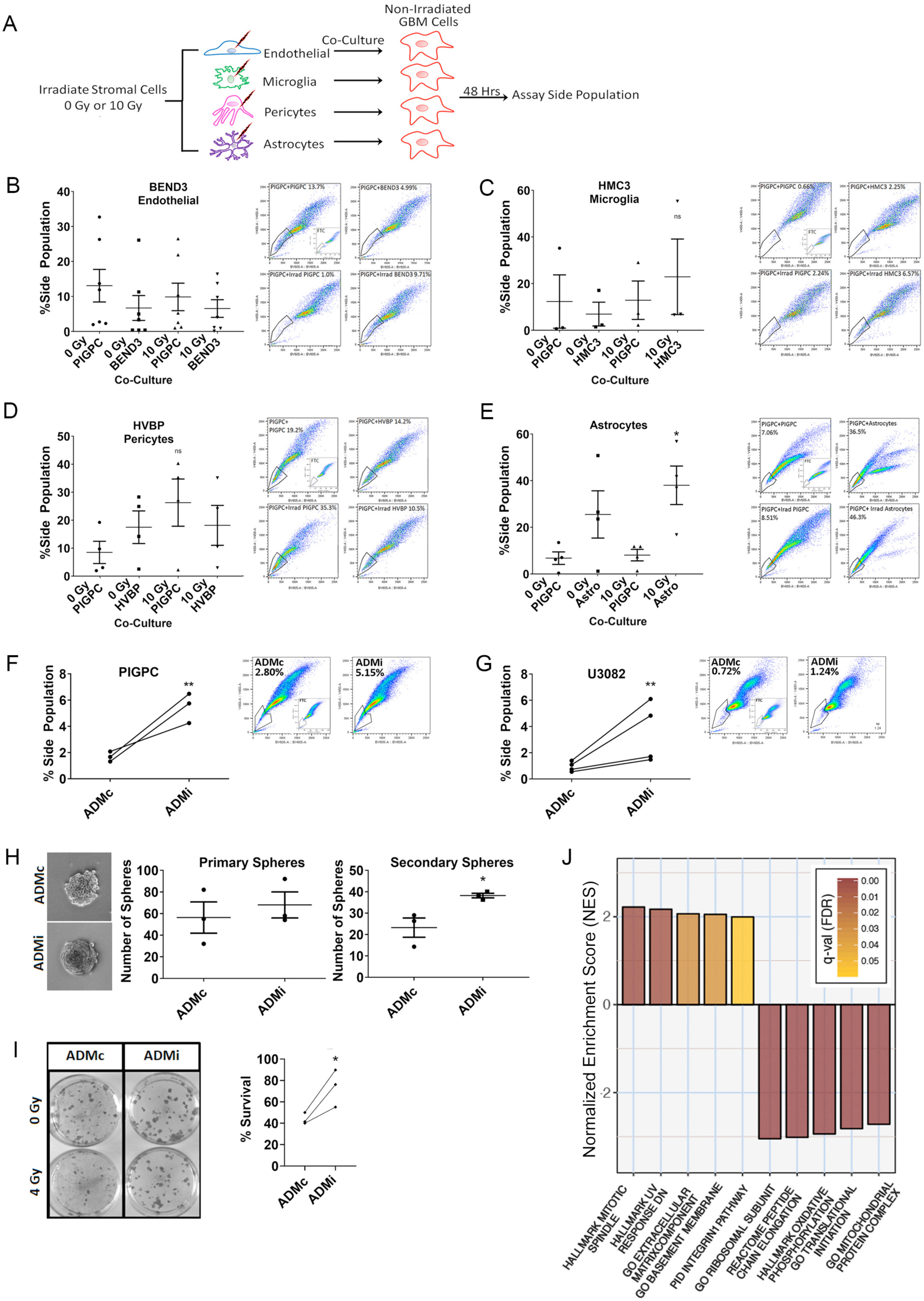
Irradiated astrocytes modify the extracellular matrix to promote glioma cell stemness and radiation resistance. **(A)** Experimental design. (**B-E**) SP of PIGPC after co-culture with the indicated cell type pre-treated with the indicated radiation dose. Dots represent individual experiments and bars represents average of all experiments with SEM. Sample FACS plots show gating for SP. (**B**) SP of PIGPC after co-culture with either 0 Gy-treated PIGPC (0 Gy PIGPC), 0 Gy-treated BEND3 endothelial cells (0 Gy BEND3), 10 Gy-treated PIGPC cells (10 Gy PIGPC), or 10 Gy-treated BEND3 cells (10 Gy BEND3). (**C**) SP of PIGPC after co-culture with either 0 Gy PIGPC, 0 Gy-treated HMC3 microglia (0 Gy HMC3), 10 Gy PIGPC, or 10 Gy-treated HMC3 cells (10 Gy HMC3). (**D**) SP of PIGPC after co-culture with either 0 Gy PIGPC, 0 Gy-treated HVBP pericytes (0 Gy HVBP), 10 Gy PIGPC, or 10 Gy-treated HVBP cells (10 Gy HVBP). (**E**) SP of PIGPC after co-culture with either 0 Gy PIGPC, 0 Gy-treated astrocytes (0 Gy Astro), 10 Gy PIGPC, or 10 Gy-treated astrocytes (10 Gy Astro). * P ≤ 0.05 by one-way ANOVA with Dunnet’s multiple comparison’s post-test. (**F**) Scatter plot showing percent SP of PIGPC cultured on ADMc or ADMi from n=3 experiments and sample plots showing an example of gating for SP. Error bars are SEM. ** P ≤ 0.01, unpaired t-test. (**G**) Scatter plot showing SP of U3082 glioma cells cultured on ADMc or ADMi in serum-free conditions (n=4) and sample plots showing an example of gating for SP. Error bars are SEM. ** P ≤ 0.01, ratio paired t-test. (**H**) Self-renewal of U3082 glioma cells pre-conditioned on ADMc or ADMi. Quantification of primary spheres (center) and secondary spheres (right) (n=3) with images of sample secondary spheres from the indicated pre-conditioning (left). Error bars are SEM. * P ≤ 0.05, unpaired t-test. (**I**) Clonal survival of U251 glioma cells plated on ADMc or ADMi in triplicate wells, followed by irradiation with 0 Gy or 4 Gy. Scatter plot indicates percent survival (n=3 independent experiments). Example colonies are shown below. Error bars are SEM. * P ≤ 0.05, unpaired t-test. (**J**) Sample of most altered gene sets in U3082 cells cultured on ADMi compared to ADMc. NES indicates change in the enrichment score of cells cultured on ADMi relative to ADMc.

We pre-treated mouse brain endothelial cells (BEND3; Fig 1B), human microglia (HMC3; Fig 1C), primary human brain vascular pericytes (HBVP; Fig 1D), and primary human astrocytes (Fig 1E) with 0 Gy or 10 Gy, which is a therapeutic radiation dose in our tumor models. PIGPC were used as control, non-stromal cells in each experiment to determine if co-culture with irradiated tumor cells alone could promote stemness features and to control for cell density. Only co-culture with pre-irradiated astrocytes stimulated a marked and consistent increase in SP of the PIGPC cells (Fig 1E), suggesting that irradiated astrocytes support glioma cell stemness.

To rule out that stromal cells alone might contribute to the SP, we measured the SP of all the cell types. Neither BEND3, HBVP, HMC3, nor astrocytes alone had a consistent SP before or after irradiation (Fig S1C-D). Co-culture of PIGPC with irradiated PIGPC also did not result in a significant increase in SP in any of the co-culture assays (Fig 1B-E).

Co-culture of astrocytes with glioma cells may increase the glioma cell SP through direct cell-cell contact, soluble factors, or modification of the extracellular matrix (ECM). We devised two assays to isolate the effects of soluble factors and ECM components from irradiated astrocytes on glioma cells (Fig S2A). To determine whether pre-irradiated astrocytes stimulated increased SP of PIGPC via soluble factors, we created conditioned media by suspending astrocytes in sodium alginate beads followed by treatment with 0 Gy or 10 Gy, then co-culture with PIGPC adhered to culture dishes. We did not detect an increase in SP of PIGPC cells cultured in this conditioned media (Fig S2B-C), suggesting the possibility of insoluble factors influencing the SP of the PIGPC cells. We therefore tested the potential of matrix proteins to affect the SP.

To generate purified ECM, confluent astrocytes were treated with 0 Gy or 10 Gy, cultured for 10 days, then de-cellularized, leaving behind insoluble matrix proteins (astrocyte derived matrix, ADM). Glioma cells were cultured on matrix from 10 Gy-irradiated astrocytes (ADMi) or control, 0 Gy-treated astrocytes (ADMc) to examine the influence of ADM in various assays measuring stemness and radiation resistance including SP, self-renewal, and survival after irradiation (Fig S2A).

Both PIGPC (Fig 1F) and the serum-free cultured human glioma cell line U3082 (Fig 1G) showed increased SP when cultured on ADMi. This suggests that changes in astrocyte-derived matrix proteins after irradiation may be responsible for the increased SP of glioma cells co-cultured with irradiated astrocytes (Fig 1E). Another important measure of stemness is self-renewal capabilities, which can be measured by sphere formation capacity at clonal densities. To determine whether ADMi affected self-renewal, U3082 cells were conditioned by culturing for 5 days on ADMc or ADMi, followed by plating at clonal densities in triplicate in 6-well dishes. After initial sphere formation, spheres were dissociated and single cells were replated at clonal densities. There were no changes in primary spheres numbers (Fig 1H), however quantification of secondary spheres revealed increased sphere formation from cells cultured on ADMi compared to ADMc (Fig 1H). The increase in SP and self-renewal of glioma cells cultured on ADMi compared to those cultured on ADMc suggests that astrocytes modify the ECM after irradiation to support glioma cell stemness.

To determine whether ADMi promotes radiation resistance of glioma cells, we plated clonal densities of human glioma U251 cells on ADMc or ADMi, followed by irradiation at 4 Gy, a dose that reduced colony formation by approximately 50% (Fig 1I). Glioma cells cultured on ADMi showed increased colony formation after irradiation compared to cells cultured on ADMc (Fig 1I). This suggests that matrix from irradiated astrocytes has the capacity to promote radiation resistance of glioma cells.

RNA sequencing of U3082 cells cultured on ADMc or ADMi for 48 h revealed that culture on ADMi resulted in global gene expression changes of glioma cells. Among gene sets significantly upregulated after culture on ADMi were pathways associated with the ECM, basement membrane, and integrin signaling (Fig 1J). Notable gene sets among the downregulated ones were associated with translation initiation and oxidative phosphorylation (Fig 1J).

Taken together, these data suggest that astrocytes modify matrix proteins after irradiation in a manner promoting stemness, including radiation resistance of glioma cells.

### The irradiated brain microenvironment supports tumor growth

Finding that tumor-naïve astrocytes irradiated in culture can promote the SP of glioma cells, we sought to determine whether astrocytes in the tumor-naïve brain might show increases in activation in response to irradiation. We stained brains from tumor-naïve mice treated with 0 Gy or 10 Gy for GFAP, one of the markers of astrocyte activation, and identified a trend of increased GFAP in the brains of 10 Gy-treated mice (Fig 2A-E). This is consistent with previous findings at higher radiation doses (Chiang et al. 1993).

**Figure 2.**
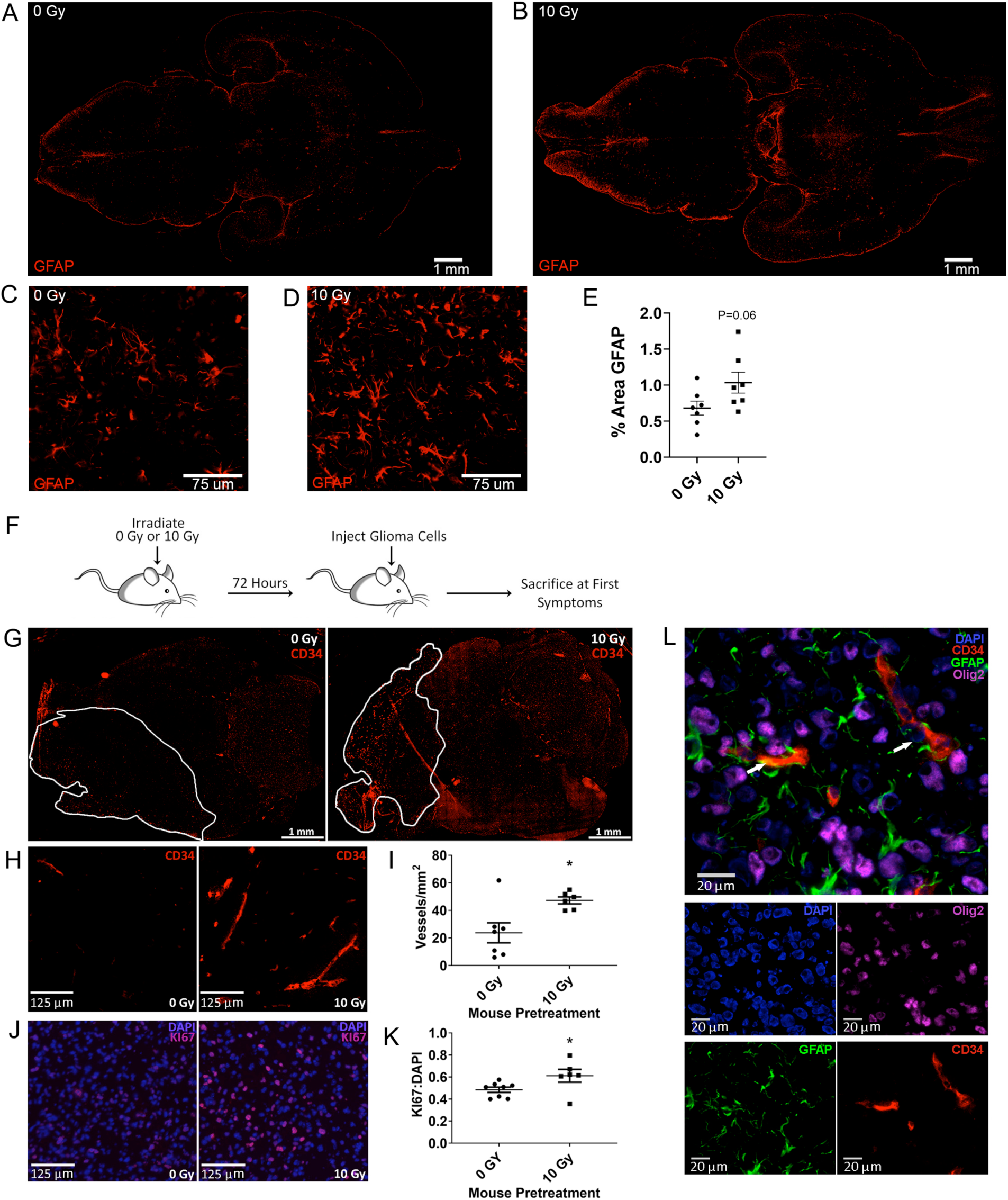
The irradiated brain microenvironment supports tumor growth. (**A-B**) Representative immunofluorescent detection of GFAP on whole brain scans of sections from non-tumor bearing mice treated with **(A)** 0 Gy or **(B)** 10 Gy. (**C-D**) High magnification image of GFAP-positive cells from A-B. (**E**) Percent area GFAP staining in 0 Gy and 10 Gy treated mice. Dots represent individual mice and bars average of all mice with SEM. P value, unpaired t-test. (**F**) Experimental design. (**G**) Whole brain scans of sections from mice pre-treated with the indicated dose of radiation prior to injection with glioma cells. Tumor-bearing brains stained for CD34 to identify vasculature. (**H**) Examples of CD34 immunofluorescence marking microvascular proliferation in mouse gliomas from mice pre-treated with the indicated dose of radiation. (**I**) Quantification of microvascular proliferation in mice pre-treated with 0 Gy or 10 Gy. (**J**) Examples of Ki67 (magenta) staining of tumors from mice pre-treated with the indicated dose of radiation and (**K**) quantification of Ki67 relative to DAPI in those tumors. Dots represent results from individual mice. (**I**,**K**) Bars represent average from n=8 0 Gy control mice and n=6 10 Gy irradiated mice. Error bars are SEM. * P ≤ 0.05, unpaired t-test. (**L**) Example of immunofluorescent staining of CD34 (red), glioma cells (Olig2, magenta), and the astrocyte marker GFAP (green). DAPI stains indicate nuclei (blue). White arrow indicates locations of association between astrocytes and vasculature.

After finding that pre-irradiated astrocytes could promote stemness features of glioma cells and irradiation alone induces reactive gliosis in tumor-naïve brains, we tested whether radiation-induced changes in the brain microenvironment influence tumor growth or development. We therefore pre-irradiated mice with 0 Gy or 10 Gy to the brain, followed by intracranial injection with PIGPC 72 h post-irradiation. Mice were sacrificed upon development of glioma symptoms (Fig 2F). Pre-irradiated mice developed tumors with increased microvascular proliferation (Fig 2G-I) and increased Ki67 positive nuclei (Fig 2J-K), both indicators of more aggressive, high grade glioma. Astrocytes were abundant within the tumor and associated with sites of microvascular proliferation (Fig 2L), one well established site enriched with stem-like glioma cells (Calabrese et al. 2007). These findings suggest that irradiation of the brain microenvironment supports tumor growth and that astrocytes may play a role in this process.

### Irradiation induces a reactive astrocyte phenotype, which persists within the original tumor volume after irradiation

Astrocytes are an abundant cell type in the brain and in gliomas and are likely to survive radiotherapy (Schneider et al. 2012). It is well established that astrocytes respond to damage to the brain (Sofroniew and Vinters 2010), including radiation,(Chiang et al. 1993; Noel and Tofilon 1998) and that this response typically involves a process termed reactive gliosis. This response varies across regions of the brain and damage type; however, some hallmarks of reactive gliosis are commonly seen (Sofroniew and Vinters 2010). including GFAP and vimentin upregulation and morphological reorganization (Sofroniew and Vinters 2010). Notably, tumors frequently recur in the astrogliotic regions overlapping with previous irradiation fields (Lemee et al. 2015).

To understand the response of astrocytes to radiation in our models, we examined both purified human astrocytes and astrocytes from normal and tumor-bearing mouse brains. Tumors were generated in *Nestin-tv-a Ink4a/Arf*^*-/-*^ mice by injection with RCAS-*PDGFB* and were irradiated upon symptoms of glioma with 0 Gy or 10 Gy, then sacrificed 72 h later (Fig 3A). Whole brain sections were stained for Olig2 (green), which marks glioma cells, and GFAP (red), which marks activated astrocytes (Fig 3B-C). In 0 Gy-treated tumor-bearing brains, reactive astrocytes are enriched at the tumor borders and are found throughout the tumor itself, as can be seen in scans of the whole brain sections (Fig 3B). Tumor area is demarcated with a white line. Closer examination reveals astrocytes found within stromal compartments (white arrow in high-magnification, merged image) and amongst the Olig2-positive glioma cells (blue arrow) (Fig 3B).

**Figure 3.**
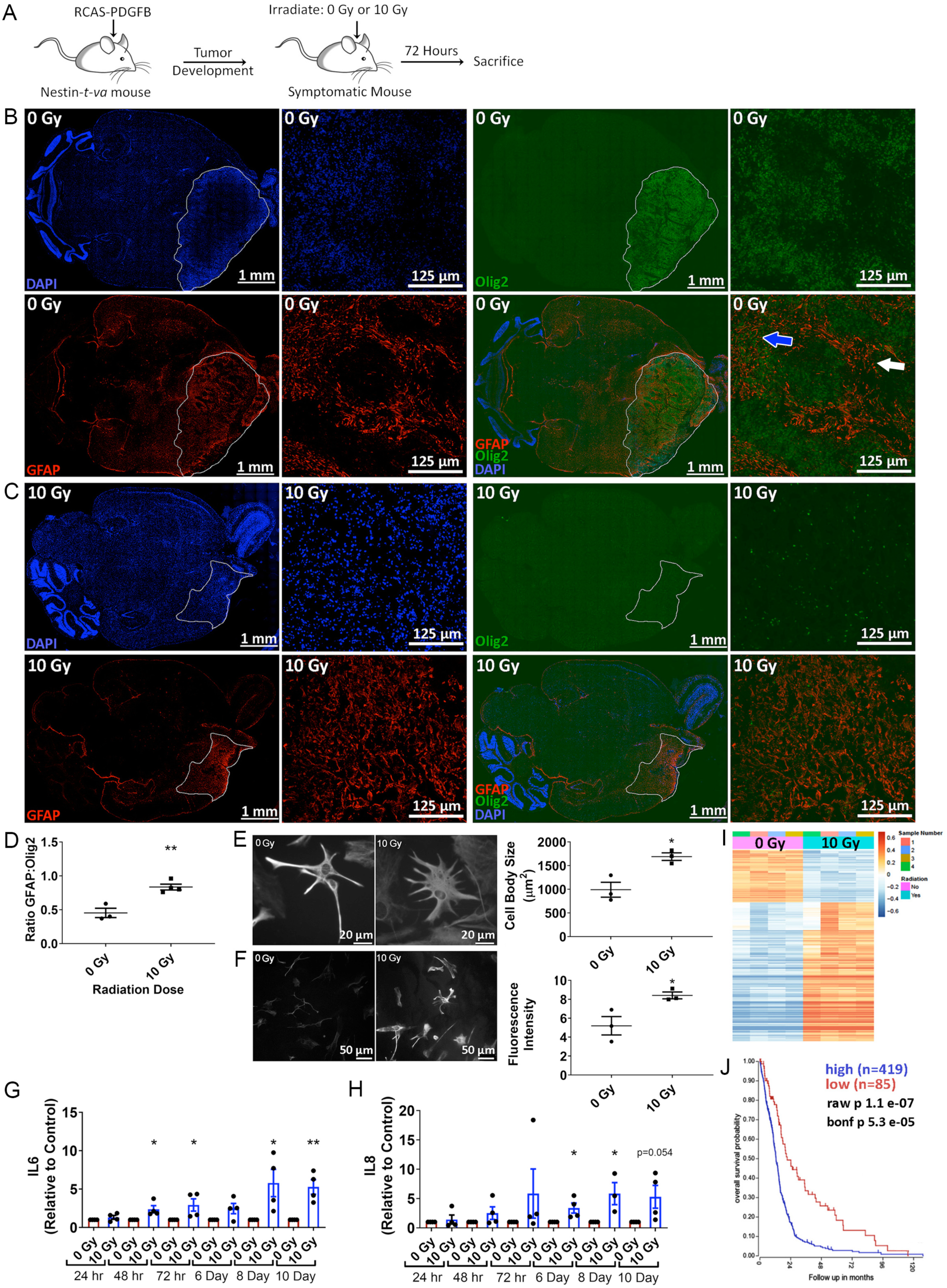
Irradiation induces a reactive astrocyte phenotype which persists within the original tumor volume after irradiation. (**A**) Experimental design. (**B-C**) Sections of brains from Nestin-*tv-a* mice injected with RCAS-*PDGFB* and treated upon symptoms with (**B**) 0 Gy or (**C**) 10 Gy. Immunofluorescent detection of astrocytes with GFAP (red) and tumor with Olig2 (green) in whole brain scans and high magnification details from the tumor region of mouse brain sections. Nuclei are stained with DAPI (blue). White outline in B indicates Olig2-positive tumor area. White line in C indicates probable tumor location before irradiation based on site of injection and GFAP-positivity. (**D**) Quantification of GFAP relative to Olig2 measured in B (0 Gy) within Olig2-positive tumor area or C (10 Gy) within area of retained GFAP positivity. White line in C indicates example of boundary where GFAP:Olig2 ratio was measured. Scatter plot of n=3 mice from each treatment. * P ≤ 0.05, unpaired t-test. (**E**) Immunofluorescent detection of GFAP in cultured astrocytes 24 h after 0 Gy or 10 Gy and quantification of astrocyte body size (n=3). * P ≤ 0.05 unpaired t-test. (**F**) Immunofluorescent detection of vimentin in cultured astrocytes 24 h after 0 Gy or 10 Gy and quantification of fluorescence intensity normalized to number of nuclei in image (n=3). * P ≤ 0.05, ratio paired t-test. (**G-H**) Quantification by ELISA of (**G**) IL6 and (**H**) IL8 in conditioned medium from astrocytes the indicated time after treatment with 0 Gy or 10 Gy (n=3). * P ≤ 0.05, ** P ≤ 0.01, unpaired t-test. ELISA results were normalized to total protein. Scatter plots represent values relative to the respective time point control. (**I**) Heatmap of RNA sequencing results of astrocytes 24 h after 0 Gy or 10 Gy. (**J**) Kaplan-Meier curve showing survival of glioma patients with high (blue) or low (red) expression of genes expressed by irradiated astrocytes. All error bars are SEM.

After irradiation, Olig2-positive cells decrease to nearly undetectable levels; however, GFAP staining is retained in areas of the likely original tumor volume (Fig 3C, white borders, identified by GFAP staining). In the areas demarcated by strong GFAP staining (white borders), this results in an increased ratio of GFAP to Olig2 in irradiated brains compared to untreated brains (Fig 3D).

Because *in vitro* cultivation of primary astrocytes shifts them to a more reactive phenotype than seen *in vivo*,(Liddelow and Barres 2017) we tested whether our primary cultured astrocytes underwent any features of reactive gliosis. Astrocytes exposed to 10 Gy developed the classic morphology of somatic hypertrophy (Fig 3E) and expressed elevated levels of the reactive marker vimentin (Fig 3F) (Liddelow and Barres 2017). Furthermore, irradiated astrocytes expressed increased levels of the soluble proteins IL6 (Fig 3G) and IL8 (Fig 3H), a response that remained for up to ten days, indicating a persistant response to the intial irradiation.

To further characterize the astrocytic response to radiation in culture, we performed RNA sequencing on primary astrocytes 24 h after receiving a dose of 0 Gy or 10 Gy, which revealed distinct gene expression patterns that differed between control and irradiated astrocytes (Fig 3I), including an upregulation of the reactive astrocyte marker GFAP (p=4.6E-16). Analysis of gene set enrichment data revealed an atypical response to irradiation, including suppression of the p53 pathway (Table S1), which has been seen previously.(Gong et al. 2017) Other notable changes to gene sets were those involved in DNA repair and cell cycle-related gene sets (Table S1). This fits with previous reports that astrocytes respond to radiation by undergoing reactive gliosis, a state in which they can become proliferative, and supports the potential for astrocytes to survive irradiation and persist in the irradiated tumor volume where they might modify the microenvironment.

Interestingly, a gene signature consisting of the 100 most upregulated genes in irradiated astrocytes as compared to controls was significantly associated with worse survival in GBM, based on analysis of The Cancer Genome Atlas (TCGA) GBM dataset (2008) (Fig 3J).

### Irradiated astrocytes secrete TGM2 *in vitro* and *in vivo* after irradiation

To identify which proteins in the ECM might be responsible for increased stemness and radioresistance of cells cultured on ADMi, we performed mass LC-MS/MS analysis on ADMc and ADMi (Fig 4A). Table S2 lists proteins that changed significantly in matrix from irradiated astrocytes (summarized graphically in Figure 4A). From amongst the potential targets identified within this altered ECM, we selected TGM2 for further inquiry because of its proposed roles in matrix modification and stemness (Bagatur et al. 2018; Condello et al. 2018), as well as promising results in a previous glioma xenograft model (Yin et al. 2017) and reports that it is a prognostic indicator of shorter time to relapse after chemoradiotherapy (Deininger et al. 2000).

**Figure 4.**
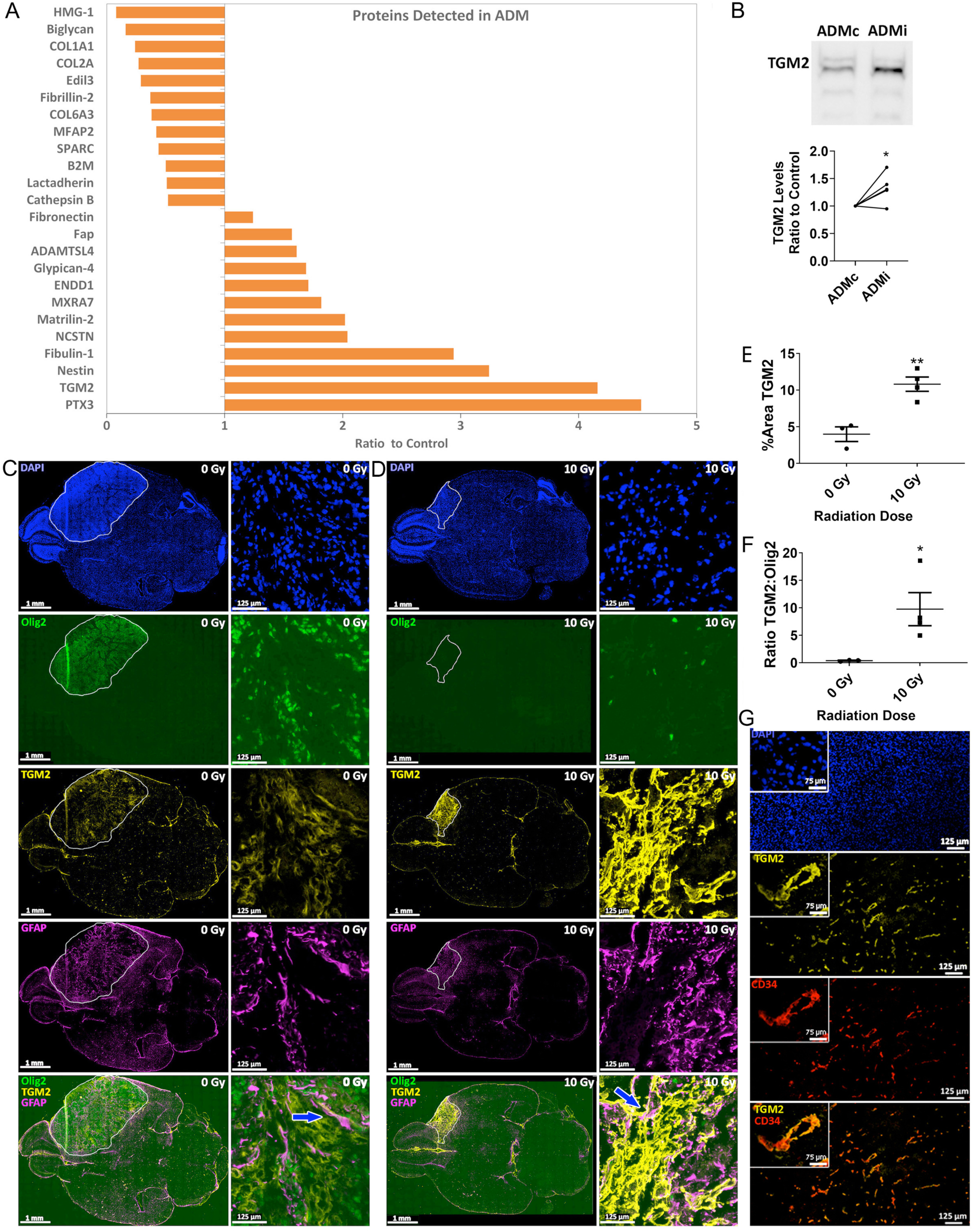
Irradiated astrocytes secrete TGM2 *in vitro* and *in vivo* after irradiation. (**A**) Mass spectrometry results of proteins detected in ADMi relative to ADMc. Matrix proteins with the largest and most significant increase or decrease in ADMi relative to ADMc. (**B**) Sample western blot of TGM2 in ADMc and ADMc. Quantification from n=6 independent samples of ADMc and ADMi. Error bars are SEM. * P ≤ 0.05 by ratio paired t-test. (**C-G**) Sections of brains from Nestin-*tv-a* mice injected with RCAS-*PDGFB* and treated upon symptoms with either (**C**) 0 Gy or (**D**) 10 Gy. Immunofluorescent detection with Olig2 (green, tumor), TGM2 (yellow), and GFAP (magenta, astrocytes) in whole brain scans and high magnification details of mouse brain sections are presented. Nuclei are stained with DAPI (blue). White outline in C indicates Olig2-positive tumor area. White line in D indicates probable tumor location before irradiation based on site of injection and TGM2-positivity. (**E**) Quantification of percent area covered by TGM2 within the non-irradiated Olig2-positive tumor area (white outline in C or within the area of retained TGM2 positivity in the irradiated tumors. White line in D indicates example of boundary where TGM2 was measured. Scatter plot represents n=3 mice from each treatment. Error bars are SEM. * P ≤ 0.05, unpaired t-test. (**F**) Quantification of TGM2 relative to Olig2 measured in C (0 Gy) within Olig2-positive tumor area or D (10 Gy) within area of retained TGM2 positivity. White line in D indicates example of boundary where TGM2 to Olig2 ratio was measured. Scatter plot represents n=3 mice from 0 Gy and n=4 mice from 10 Gy treatment. Error bars are SEM. * P ≤ 0.05, unpaired t-test. (**G**) Example of CD34-labeled blood vessels (red) co-stained with TGM2 (yellow) with high magnification inset.

We first confirmed increased expression of TGM2 in matrix from multiple batches of astrocytes from different individuals by western blot (Fig 4B). We followed this by examining TGM2 expression *in vivo* in our genetic model of PDGFB-driven gliomagenesis (Fig 3A). Whole brain sections were stained for Olig2 (green), TGM2 (yellow), and GFAP (magenta) (Fig 4C-D). Relative to non-tumor bearing brain tissue, TGM2 expression was elevated in tumors (white bounded area, Fig 4C), was extensively expressed in the stromal compartment, and showed areas of co-staining with GFAP (Fig 4C, example: blue arrow, merged high magnification image). In the tumors treated with 10 Gy (Fig 4D), TGM2 expression was retained in the likely original tumor volume and even increased, resulting in a greater percent area covered by TGM2 (Fig 4E) and increased ratio of TGM2 to Olig2 (Fig 4F) within the area of TGM2-positive staining compared to the non-irradiated brain. In the irradiated tumors, regions of TGM2 staining were also found in close association with GFAP-positive astrocytes, often co-staining or showing astrocytes surrounding areas of intense TGM2-positivity (Fig 4D, example: blue arrow, merged high magnification image).

Another notable expression pattern for TGM2 demonstrates that, like GFAP, TGM2 expression is elevated in areas of microvascular proliferation and the normal the vasculature (Fig 4G). Comparable to our findings in mice, analysis of the Ivy Glioblastoma Atlas Project (IVY-GAP) (Puchalski et al. 2018) reveals that TGM2 is highly elevated in areas of microvascular proliferation (Fig S3).

Taken together, these data suggest that irradiation promotes increased expression of TGM2 from astrocytes, potentially providing a radio-protective environment for a newly expanding tumor. We therefore returned to our *in vitro* experimental systems to determine if there was a functional relationship between TGM2 and stemness and radiation resistance of glioma cells.

### Astrocyte-derived TGM2 promotes glioma stemness after irradiation

To determine whether TGM2 expression might be associated with glioma cell stemness or radiation resistance, we examined the effects of either purified TGM2 or TGM2 inhibitors on our *in vitro* model of glioma cell-astrocyte matrix interaction. Purified TGM2, when coated onto ADMc, raised the SP of PIGPC near to that seen with PIGPC cultured on ADMi (Fig 5A). In contrast, two different TGM2 inhibitors, GK921 (Fig 5B) and dansylcadaverine (DC; Fig 5C), decreased the SP of PIGPC cultured on ADMi to the levels seen on ADMc. GK921 also decreased survival after irradiation of PIGPC cultured on ADMi to levels seen on ADMc, while not affecting PIGPC cultured on ADMc (Fig 5D). These data suggest that TGM2 contributes to the increased stemness and radiation resistance of glioma cells cultured on ADMi as compared to those cultured on ADMc.

**Figure 5.**
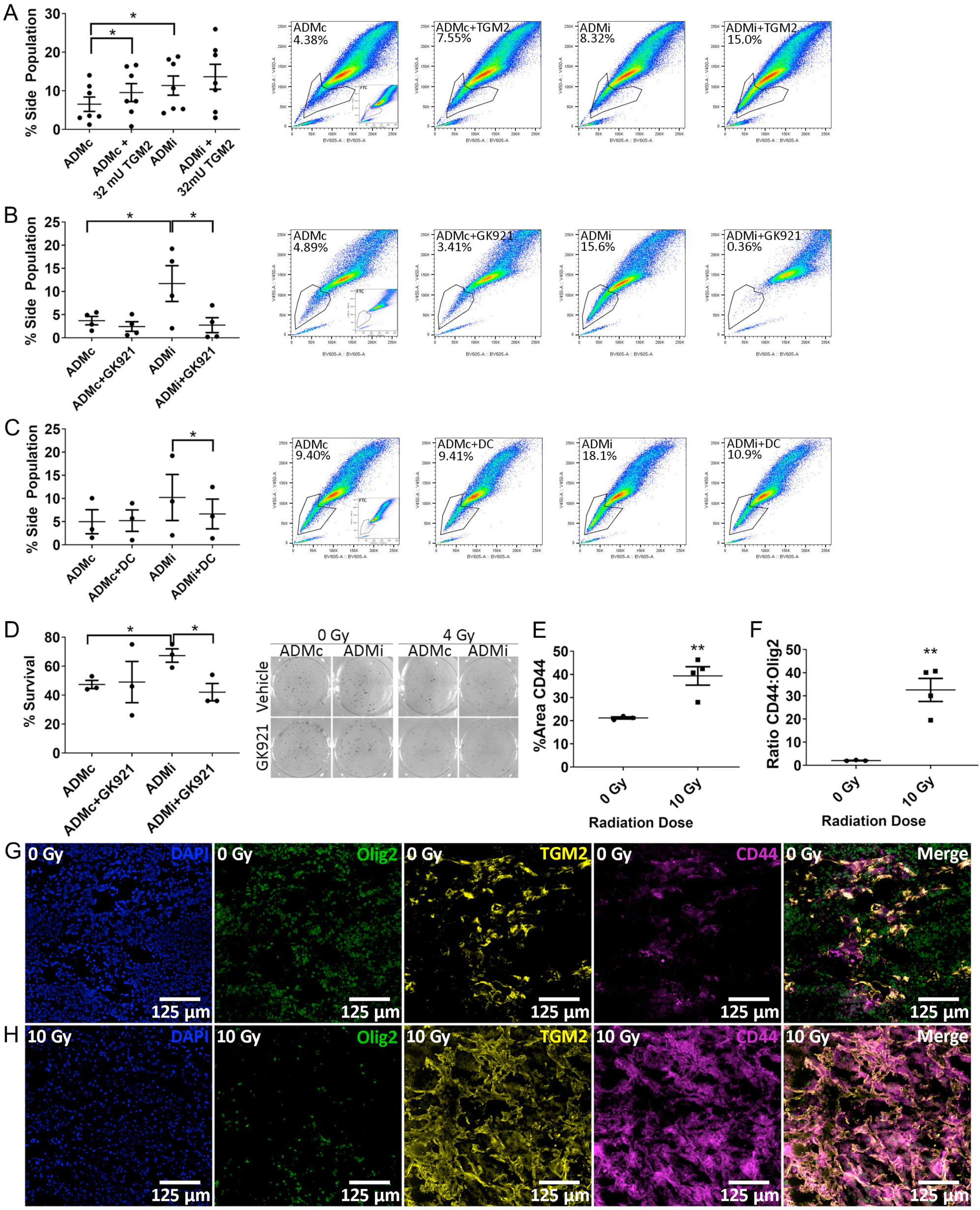
Astrocyte-derived TGM2 promotes glioma stemness after irradiation. (**A**) Quantification of n=7 independent assays in which PIGPC glioma cells were cultured on ADMc or ADMi coated with vehicle or 32 mU TGM2 (left). Sample FACS plots with gating (right) * P ≤ 0.05, one-way ANOVA with Dunnet’s multiple comparison’s post-test. (**B**) Quantification of n=4 independent assays in which PIGPC glioma cells were cultured on ADMc or ADMi with 0.5 µM of the TGM2 inhibitor GK921 (left). Sample FACs plots with gating (right). (**C**) Quantification of n=3 independent assays in which PIGPC glioma cells were cultured on ADMc or ADMi with 100 µM of the TGM2 inhibitor dansylcadaverine (DC) (left). Sample FACS plots with gating (right). (**D**) Clonal survival of U251 glioma cells plated on ADMc or ADMi in triplicate wells in media containing 0.1 µM GK921, followed by irradiation with 0 Gy or 4 Gy. Scatter plot indicates average percent survival (n=3) (left) and example colonies are shown on the right. Error bars are SEM. * P ≤ 0.05, unpaired t-test. (**E**) Quantification of percent area covered by CD44 within the non-irradiated Olig2-positive tumor area or within the area of retained CD44 positivity in the irradiated tumors. (Boundaries indicated in Figures 4 C and D by white line.) Scatter plot represents n=3 mice from 0 Gy and n=4 mice from 10 Gy treatment. Error bars are SEM. ** P ≤ 0.01, unpaired t-test. (**F**) Quantification of CD44 relative to Olig2 measured in G (0 Gy) within Olig2-positive tumor area or H (10 Gy) within area of retained TGM2 positivity. (Boundaries indicated in Figures 4 C and D by white line.) Scatter plot represents n=3 mice from 0 Gy and n=4 mice from 10 Gy treatment. Error bars are SEM. * P ≤ 0.05, unpaired t-test.

*In vivo*, tumors treated with 10 Gy increased expression of CD44 (Fig 5E-H), a stem cell marker in glioma(Anido et al. 2010) and a marker of mesenchymal subtype GBM, which strongly co-localized with TGM2 (Fig 5G-H). Like TGM2, CD44 expression was retained in the likely original tumor volume after irradiation, even as the Olig2-positive cells receded, as seen by increased expression relative to Olig2 (Fig 5F). These data provide evidence that TGM2 is associated with increased stemness *in vivo* and is elevated after irradiation.

### TGM2 is a potential therapeutic target in GBM

To further examine the effects of TGM2 inhibition, we used an *ex vivo* organotypic slice model of GBM. We initiated tumors in our *Nestin-tv-a Ink4a/Arf*^*-/-*^ mice by injection with RCAS-*PDGFB* and shp53 and then sliced freshly-harvested brains from mice upon detection of glioma symptoms. Slices were cultured overnight, followed by treatment with 0 µM or 5 µM of the TGM2 inhibitor GK921 for 72 h, then fixed and cryosectioned, followed by immunofluorescent detection of Olig2 to determine the presence of tumor cells. TGM2 inhibitor significantly decreased the percent of Olig2-positive nuclei in the sections, in some cases reducing Olig2-positive cells to undetectable levels (Fig 6A-C).

**Figure 6.**
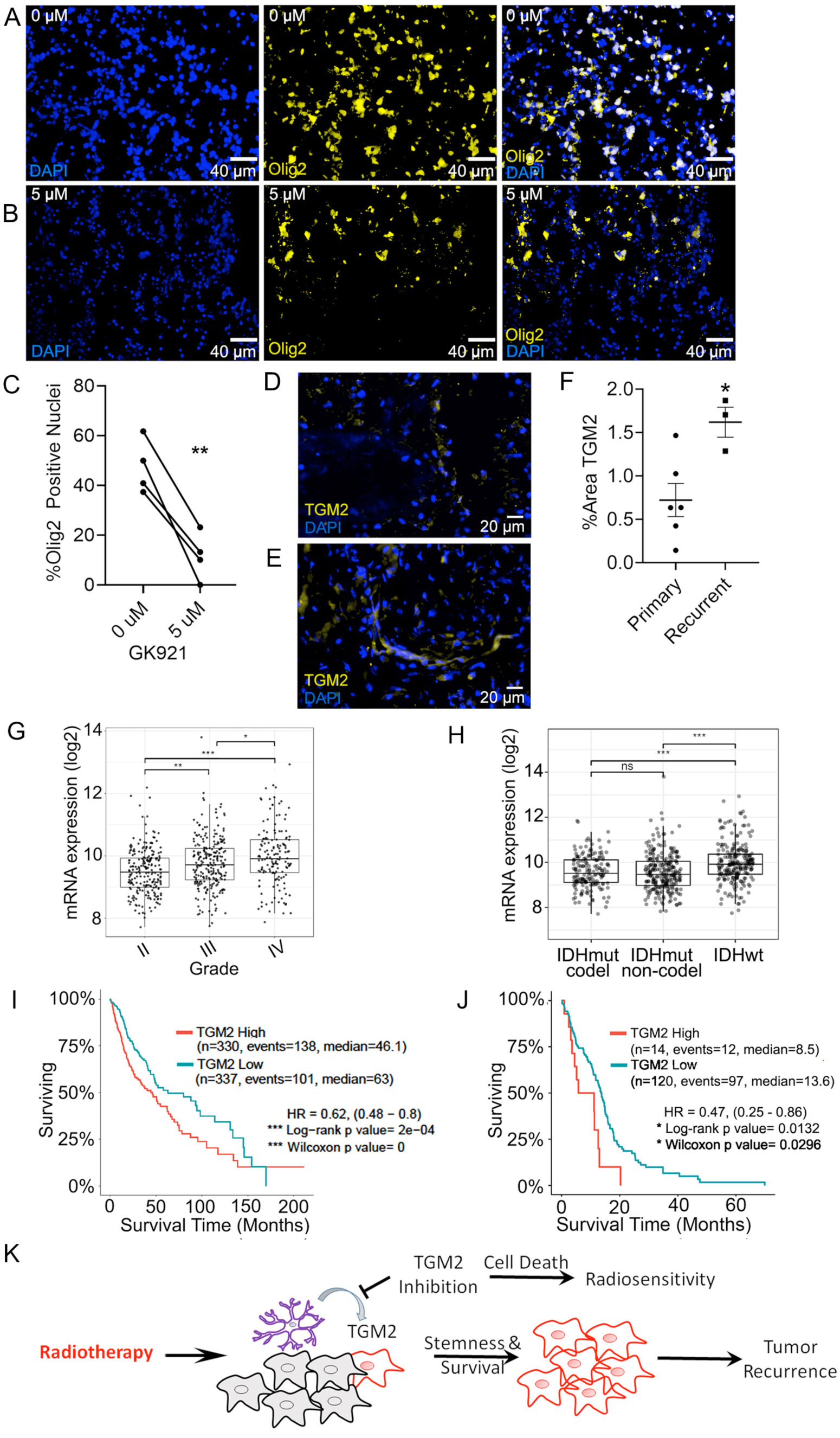
TGM2 is a potential therapeutic target in GBM. (**A-B**) Immunofluorescent detection of Olig2 (yellow) or nuclei (DAPI, blue) in cryosections of organotypic brain slices from mice bearing gliomas. Brain slices were treated with either (**A**) 0 µM or (**B**) 5 µM of the TGM2 inhibitor GK921. (**C**) Quantification percent of nuclei positive for Olig2 in organotypic brain slices from four mice. (**D-F**) Immunofluorescent detection of TGM2 in human gliomas in either (**D**) primary glioma or (**E**) recurrent glioma. (**F**) Scatter plot showing elevated TGM2 in recurrent glioma. Each point represents the average percent area of TGM2 from six areas measured within a tumor section from one of six patients (primary) or one of three patients (recurrent). * P ≤ 0.05, unpaired t-test. (**G**) TGM2 transcript expression in glioma grades II, III, and IV. H) TGM2 transcript expression in IDH mutant (IDHmut) glioma with or without 1p/19q codeletion (codel and non-codel respectively) or IDHwt glioma. (**I**) Kaplan-Meier curve showing survival of glioma patients with either high (red) or low (blue) TGM2 expression. (**J**) Kaplan-Meier curve showing survival of GBM patients with IDHwt tumors with TGM2 transcript expression in the upper 10% (red) or lower 90% (blue). (**K**) Proposed model of stimulation of TGM2 release from astrocytes after irradiation and its role in survival of tumor cells. Grey cells, glioma cells that respond to radiotherapy. Red cells, glioma cells that survive radiotherapy. Purple cells, astrocytes.

A small sample of human GBM shows elevated TGM2 protein expression in recurrent versus primary GBM (Fig 76D-F), suggesting higher TGM2 has the potential to remain in the GBM microenvironment at recurrence. All patient samples were isocitrate dehydrogenase wild type (IDHwt). Recurrent tumors were from patients receiving radiotherapy of 60 Gy and temozolomide adjuvant therapy. Additionally in human glioma, TGM2 mRNA is elevated as tumor grade increases (Fig 6G). TGM2 mRNA is significantly increased in IDHwt tumors (Fig 6H), the most aggressive tumor classification (Louis et al. 2016). Elevated TGM2 correlates with shorter survival times (Fig 6I). Within IDHwt GBM, tumors with the top 10% of TGM2 mRNA expression have the shortest survival times (Fig 6J). These data suggest that TGM2 may be a viable target in GBM.

Based on our findings, we propose a model in which radiation stimulates tumor-associated astrocytes to secrete TGM2, which then promotes stemness and survival of glioma cells, allowing tumor recurrence (Fig 6K). Our findings suggest that inhibition of TGM2 signaling provides a potential new therapeutic target to sensitize glioma cells to radiotherapy and prevent recurrence (Fig 6K).

## Discussion

GBM treatment options are limited after tumor recurrence. As surgery can rarely eliminate all GBM cells, identifying and targeting pathways that support GBM cell survival after radiotherapy is essential to improving outcomes. Clinical findings indicate that a short course of radiotherapy over three weeks results in no clinically significant difference in outcomes compared to a six-week course (Roa et al. 2004; Roa et al. 2015). One interpretation of these data, in light of findings presented here, is that neuroprotective pathways may be initiated in response to radiation to protect cells from further radiation-induced damage, and that GBM cells can take advantage of this altered microenvironment. While studies have begun examining how the tumor microenvironment can promote radiation resistance and GBM stemness (Jamal et al. 2010; Silver and Lathia 2018; Hide et al. 2019; Li et al. 2019), little study has been made of how the stromal cells themselves respond to treatment modalities. This is an important question, as findings presented here suggest that stromal cells may respond to radiation in ways which could prime the microenvironment for protection of glioma cells against further radiation insults.

It is likely that effects of radiotherapy on the tumor microenvironment are not limited to its effect on astrocytes described here. While our *in vitro* co-culture screen of pre-irradiated stromal cells detected a SP increase only from pre-irradiated astrocytes, our screen may favor our ability to detect the effect of irradiated astrocytes on glioma cells over other cell types. Furthermore, the use of the SP assay as a readout in our screen limits our findings to alterations that affect this one property of glioma cells. Nevertheless, this screen resulted in the identification of a potential new target for glioma therapy. Screens performed in other model systems may reveal further important interaction among pre-irradiated stromal cells and glioma cells.

This study provides a unique and promising new strategy in GBM treatment by examining lasting changes to the tumor microenvironment made by stromal cells in response to radiotherapy, changes which may support radiation resistance of GBM cells. While previous work has shown that in untreated tumors, astrocytes react to the GBM microenvironment and can promote glioma growth (Mega et al. 2020), we have identified astrocytes as persisting within the original tumor volume even after the tumor bulk is killed by radiation, altering that environment in a radioprotective manner. We demonstrate that they do this by secreting TGM2, which supports stemness and radiation resistance of GBM cells, which may further elucidate previous findings that TGM2 inhibition inceased survival in a glioma xenograft model (Yin et al. 2017).

Although we identified TGM2 as a potential mediator of radiation resistance and increased stemness, the mechanism by which TGM2 protects GBM cells is as yet unclear. However, fibronectin bundling, which is perhaps the most well described function of TGM2, influences radiation resistance (Serres et al. 2014), angiogensis (Giannopoulou et al. 2001), and invasiveness (Cordes et al. 2003) in many cancers, and may generate these effects through integrinβ1 signaling (Cordes et al. 2003; Cordes et al. 2006; Lugano et al. 2018) or p53 suppressive pathways (Yu et al. 2018). Which of these or other pathways mediate the pro-survival effects of TGM2 warrants further study. Our findings suggest a new strategy for sensitizing GBM cells to radiotherapy by short-circuiting the normal brain’s protective response to radiation.

## Materials and Methods

### Generation of Murine Gliomas

Gliomas were induced in Nestin-*tv-a* or Nestin-*tv-a Ink4a/Arf*^-/-^ mice by injecting indicated combinations of RCAS-*PDGFB* and RCAS-shp53-transfected DF-1 cells (ATCC) intracranially in the brain as previously described (Holland et al. 1998; Ozawa et al. 2014). Mice were monitored daily and euthanized upon exhibiting glioma symptoms. Sample numbers were selected to detect response above variability. All images from mouse studies were blinded for analysis.

### Cell Lines and Cultures

PIGPC were isolated as previously described (Pietras et al. 2014). PIGPC and U251 cells were cultured in DMEM (Corning) supplemented with 10% FBS (Biological Industries) and 1% PenStrep (Corning). HBVP (ScienCell) were cultured in PM (ScienCell) on 2 µg/cm^2^ poly-l-lysine mol wt 70,000-150,000 (Sigma) coated plastic and used below passage 4. HMC3 (ATCC, CRL-3304) were cultured in DMEM (Corning) supplemented with 10% FBS (Biological Industries) and 1% PenStrep (Corning). BEND3 (ATCC, CRL-2299) were cultured in Endothelial Cell Growth Medium MV2 (PromoCell) on 0.1% gelatin coated plastic. Primary human astrocytes (3H Biomedical) were cultured in Astrocyte Medium (3H Biomedical) and used below passage 15. U3082 glioma cells were obtained from HGCC (hgcc.se) and were cultured in HGC medium as previously described(Xie et al. 2015) as neurospheres or monolayer on laminin-coated (BioLamina, LN521) plastic.

### Immunofluorescence

Cryosections were fixed in 4% PFA followed by permeabilization in 0.3% Triton-x-100 and blocking in 1% BSA in PBS or DACO blocking reagent. Cryosections of organotypic slice cultures were fixed in 4% PFA followed by permeabilization in 0.1% Triton-x-100 and 0.1% sodium citrate then overnight incubation in 1% BSA or DAKO Diluent with the following primary antibodies: Fig. 2A-D: Rabbit anti-GFAP (DAKO, Z033401-2); Fig. 2G-H: Rat anti-CD34 (eBiosciece, 14-0341-81); Fig. 2J: Rabbit anti-Ki67 (ThermoFischer, 12683697); Fig. 2L: Rat anti-CD34 (eBiosciece, 14-0341-81), rabbit anti-Olig2 (Merck, AB9610) and chicken anti-GFAP (AB4674, Abcam); Fig. 3B: Rabbit anti-Olig2 (Merck, AB9610) and chicken anti-GFAP (AB4674, Abcam); Fig. 3E: Rabbit anti-GFAP (DAKO, Z033401-2); Fig. 3F: Rabbit anti-Vimentin (Abcam, AB45939); Fig. 4C: Rabbit anti-Olig2 (Merck, AB9610), sheep anti-TGM2 (Novus Biologicals, AF5418), and rat anti-GFAP (Life Technologies, 130300); Fig. 4G: Sheep anti-TGM2 (Novus Biologicals, AF5418) and Rat anti-CD34 (eBiosciece, 14-0341-81); Fig. 5G-H: Rabbit anti-Olig2 (Merck, AB9610), sheep anti-TGM2 (Novus Biologicals, AF5418), and rat anti-GFAP (Life Technologies, 130300); Fig. 6A-B: Goat anti-Olig2 (Novus, AF2418); Fig. 6D: Rabbit anti-TGM2 (Novus Biologicals, NBP1-86951).

After washing, sections were incubated with the appropriate Alexafluor-conjugated secondary antibodies for 1 h: Donkey anti-sheep 647 (Abcam, AB150179), donkey anti-rat 568 (Abcam, AB175475), donkey anti-rat 488 (Invitrogen, 10123952), donkey anti-rabbit 488 (Invitrogen, A21206), goat anti-chicken 568 (Invitrogen, 10462822). Images were acquired using an Olympus BX63 microscope and DP80 camera and cellSens Dimension v 1.12 software (Olympus).

### Co-culture experiments

For co-culture experiments 100,000 BEND3, HMC3 or PIGPC or 50,000 HBPV cells were pretreated with or without 10 Gy followed by 48h of co-culture with 100,000 PIGPC. Co-culture of microglia, pericytes, endothelial cells, or astrocytes and PIGPC was performed in DMEM.

### Irradiation of cells

Irradiated pericytes, microglia, endothelial cells, and astrocytes were pretreated with 10 Gy in a CellRad x-ray cell irradiator (Faxitron).

### Side Population Assay

For SP, cells were resuspended at 1,000,000 cells/ml and incubated at 37°C, 30 minutes with or without 10 µM Fumitremorgin C (Sigma), then for 90 minutes with 5 mg/ml Hoechst 33342 (Sigma) with periodic vortexing, then analyzed on a FACSVerse instrument (BD) with a 405 nm violet laser. Dual wavelength detection was performed using 448/45 (Hoechst 33342-blue) and 613/18 (Hoechst 33342-red) filters.

### Generation of Astrocyte Derived Matrix

Cofluent astrocytes were irradiated with 0 Gy or 10 Gy followed by 10 days of culture on 0.2% gelatin in astrocyte medium supplemented with 50 µg/ml L-ascorbic acid (Sigma). After 10 days, plates were decellularized in 0.4 mM NH_4_OH (Fluka), 0.5% Triton X-100 in PBS with 1 mM CaCl_2_ and 0.5 mM MgCl_2_ at 37 °C, washed with PBS containing 1 mM CaCl_2_ and 0.5 mM MgCl_2_, then treated with 10 µg/ml DNAse I (Roche) for 1 h at 37 °C. Plates were washed and stored in PBS containing 1 mM CaCl_2_ and 0.5 mM MgCl_2_ at 4 °C.

### Western Blot

Matrix was scraped into 175 µl 4X Laemmli buffer (Bio-Rad, 161-0747) containing 150 mM DTT and boiled for 10 min. Equal volumes of lysate were separated on 7.5% polyacrylamide followed by transfer to PVDF, blocking with 5% non-fat dry milk in PBS, then immunodetection using TGM2 antibody (Abcam, ab2386) followed by detection with HRP-conjugated anti-mouse secondary antibody (ThermoFisher, 31430) and ECL substrate (ThermoFisher, 34095). Blots were visualized on the LAS-3000 Imager (Fujifilm) and quantified using ImageJ.

### Sample preparation of Cell Derived Matrix

Cell-derived matrix was solubilized for LC-MS/MS analysis according to Naba *et al.*(Naba et al. 2015) Solid Phase Extraction with StageTips was performed according to the protocol of Rappsilber *et al*. (Rappsilber et al. 2007) using home packed C18 reversed-phase columns.

### LC-MS/MS analysis

Samples were reconstituted in 20µl 2% acetonitrile, 0.1% TFA, then run as triplicate injections on a Orbitrap Fusion Tribrid MS system (Thermo Scientific) equipped with a Proxeon Easy-nLC 1000 (Thermo Fisher). Injected peptides were trapped on an Acclaim PepMap C18 column (3 µm particle size, 75 µm inner diameter x 20 mm length), then gradient elution of peptides was performed on an Acclaim PepMap C18 column (2 µm particle size, 75 µm inner diameter x 250 mm length). The analytical column was coupled to the mass spectrometer using a Proxeon nanospray source. The mobile phases for LC separation were 0.1% (v/v) formic acid in LC-MS grade water (solvent A) and 0.1% (v/v) formic acid in acetonitrile (solvent B). Peptides were loaded with a constant pressure mode of solvent A onto the trapping column, then eluted via the analytical column at a constant flow of 300 nL/min. During elution, solvent B increased from 5% to 10% in 2 minutes, then to 25% in 50 minutes, then to 60% in 15 minutes, then to 90% in 5 minutes, and finally to 90% in 5 minutes. The peptides were introduced into the Orbitrap via a Stainless steel emitter 40 mm (Thermo Fisher) and a spray voltage of 2 kV was applied with a 275°C capillary temperture.

Data acquisition was carried out using a top N based data-dependent method with a 3 second cycle time. The master scan was performed in the Orbitrap in the range of 350–1350 *m/z* at a resolution of 120,000 FWHM. The filling time was set at maximum of 50 ms with limitation of 400,000 ions. Ion trap CID-MS2 was acquired using normal mode, filling time maximum 60 ms with limitation of 7000 ions, with a precursor ion isolation width of 0.7 *m/z* and a rapid Ion trap scan. Normalized collision energy was set to 27%. Only multiply charged (2^+^ to 5^+^) precursor ions were selected for MS2. The dynamic exclusion list was set to 30 s and relative mass window of 5ppm.

MS/MS spectra were searched with PEAKS (8.5) using UniProt Human database. Trypsin was used and 2 missed cleavages allowed. 15 ppm precursor and 0.5 Da fragment tolerance were used as mass tolerance. Oxidation (M) and deamidation (NQ) were treated as dynamic modification and carbamidomethylation (C) as a fixed modification. Maximum number of PTM per peptide was set to three. Matrix proteins with at least 2 unique peptides and a protein significance score > 50 were included in the results.

### Colony Assay

U251 cells were plated at 200 cells per well in a 6-well dish on ADMc or ADMi in DMEM. After 24 h, dishes were irradiated with 0 Gy or 4 Gy. Cells were cultured 2 weeks or until visible colonies formed. Colonies were fixed in 4% PFA, stained with 0.01% Crystal Violet for 1 hr, photographed on a LAS-3000 imaging system, then counted using Image J followed by visual confirmation.

### Self-Renewal Assay

U3082 cells were cultured on ADMc or ADMi 5 days in HGC Media. For primary spheres, cells were dissociated with Accutase then replated in 3 wells each of 6-well dishes at 300 cells/ml in 2 ml HGC Media and cultured until visible spheres formed, typically between 14 and 18 days. Primary spheres were visualized and counted, then dissociated and re-plated as described above. Spheres were photographed and counted on a Zeiss AX10 inverted microscope.

### RNA Sequencing Analyses

Sequencing performed at the Center for Translational Genomics, Lund University and Clinical Genomics Lund, SciLifeLab, and National Genomics Infrastructure Stockholm Node, Science for Life Laboratory. Raw count data was imported and analyzed using R statistic language (version 3.3.2). Data was rlog-normalized and combat corrected for experimental batch using DESeq2 and sva packages, respectively. DESeq differential expression analysis was performed on non-normalized raw counts. Genes with Benjamini adjusted p-values of <0.05 were considered differentially expressed. All genes ranked according to DESeq statistic (positive associated with highly expressed in irradiated astrocytes) and gene set enrichment analysis (GSEA)(Subramanian et al. 2005) was applied using the c2.all.v6.0 curated gene sets (MsidDB; broad.mit.edu/gsea/msigdb).

### Patient cohort analysis

The R2: Genomics Analysis and Visualization Platform (http://r2.amc.nl) was used to examine the Glioblasoma-TCGA-540 dataset.(2008) Astrocyte irradiation signature score was calculated using the z-score of the top 100 upregulated genes. Kaplan-Meier analysis using the scanning method was applied to the signature score using overall survival data and Bonferoni adjusted p-values of <0.05 was considered significant. Data from the Allen Institute for Brain Science IVY-GAP (http://glioblastoma.alleninstitute.org) were analyzed using the GlioVis data portal for visualization and analysis of brain tumor expression datasets (Bowman et al. 2017; Puchalski et al. 2018).

### Culture in Sodium Alginate

Dissociated human astrocytes or PIGPCs were suspended in sodium alginate (Aldrich) 4:5 with cell culture medium. The cells were then added dropwise in medium containing 0.1 M CaCl_2_ to form hydrogel beads and medium was changed to fresh AM or DMEM followed by culture at 37° C, 5% CO_2_.

### Organotypic Slice Culture

Freshly dissected brains were sliced into 300 µm slices using a 5100 mz vibrating blade tissue slicer (Campden Instruments). Slices were supported on polycarbonate tissue culture dish inserts with 0.4 µm pore size for culture at 37°C, 5% CO_2_ in HGC Medium. After overnight incubation, slices were treated with 0 µM or 5 µM of the TGM2 inhibitor GK921 daily for 72 h. Slices were PFA fixed then cryosectioned. After immunofluorescent detection of Olig2, percent Olig2-positive nuclei were quantified using Cell Profiler (Carpenter et al. 2006).

### Statistical Methods

All statistical analysis was performed in GraphPad Prism vs 8.1.2. Stastistical tests and number of replicates are indicated in the figure legends. All t-tests were two-tailed.

### Patient Glioma Samples

Human biological samples were collected according to the Swedish guidelines for use of clinical material, which includes written consent from patients and approvals by the regional ethical committee (Dnr.2018/37). IDH mutation status was determined according to clinical routine. Sample number was not predetermined but based on availability and images were blinded for analysis.

### Animal Use

Laboratory animal use was conducted in accordance with European Union directive on the subject of animal rights and approved by local ethical committees (CEIyBA (CBA PA37_2013)/ (IACUC 029-2013/001) (CNIO), M-186/14 (Lund), and 5.8.18-16350/2017 (Uppsala).

## Acknowledgments

We thank Christina Möller for her skillful technical assistance. The authors would like to acknowledge support from Science for Life Laboratory, the National Genomics Infrastructure, NGI, and Uppmax for providing assistance in massive parallel sequencing and computational infrastructure. We also wish to acknowledge the Center for Translational Genomics, Lund University and Clinical Genomics Lund, SciLifeLab Sequencing for sequencing performed in their labs. We acknowledge the Preclinical Cancer Treatment Center, SciLifeLab, Uppsala University, for technical assistance and support with radiation therapies experiments *in vivo*. Support from the Swedish National Infrastructure for Biological Mass Spectrometry is gratefully acknowledged. The results shown here are in whole or part based upon data generated by the TCGA Research Network: https://www.cancer.gov/tcga. This study was supported by grants from the Ragnar Söderberg Foundation, the Swedish Cancer Society, the Swedish Research Council, the Swedish Childhood Cancer Fund, Ollie & Elof Ericssons foundation, Jeanssons stiftelser, the Crafoord foundation, Gösta Miltons donationsfond, Stiftelsen Cancera, donations from Viveca Jeppson and Maj-Britt and Allan Johansson and the Seve Ballesteros Foundation. JB was supported by Region Skane and ALF.

The authors declare no conflicts of interest.

## Author contributions

A.P. and T.J.B. conceived of the study and experimental design. T.J.B., C.M., V.P., E.J., and T.B. designed, performed, and analyzed experiments. K.S., D.L. and E.J.P. analyzed data. V.G., J.B., and M.B. collected and provided patient samples. F.J.S., H.A., M.S., and A.P. supervised the research. A.P. and T.J.B wrote the manuscript. All authors contributed to the final manuscript.

