## Supplemental Data for "The irradiated brain microenvironment supports glioma stemness and survival via astrocyte-derived Transglutaminase 2"

Supplemental Figure 1

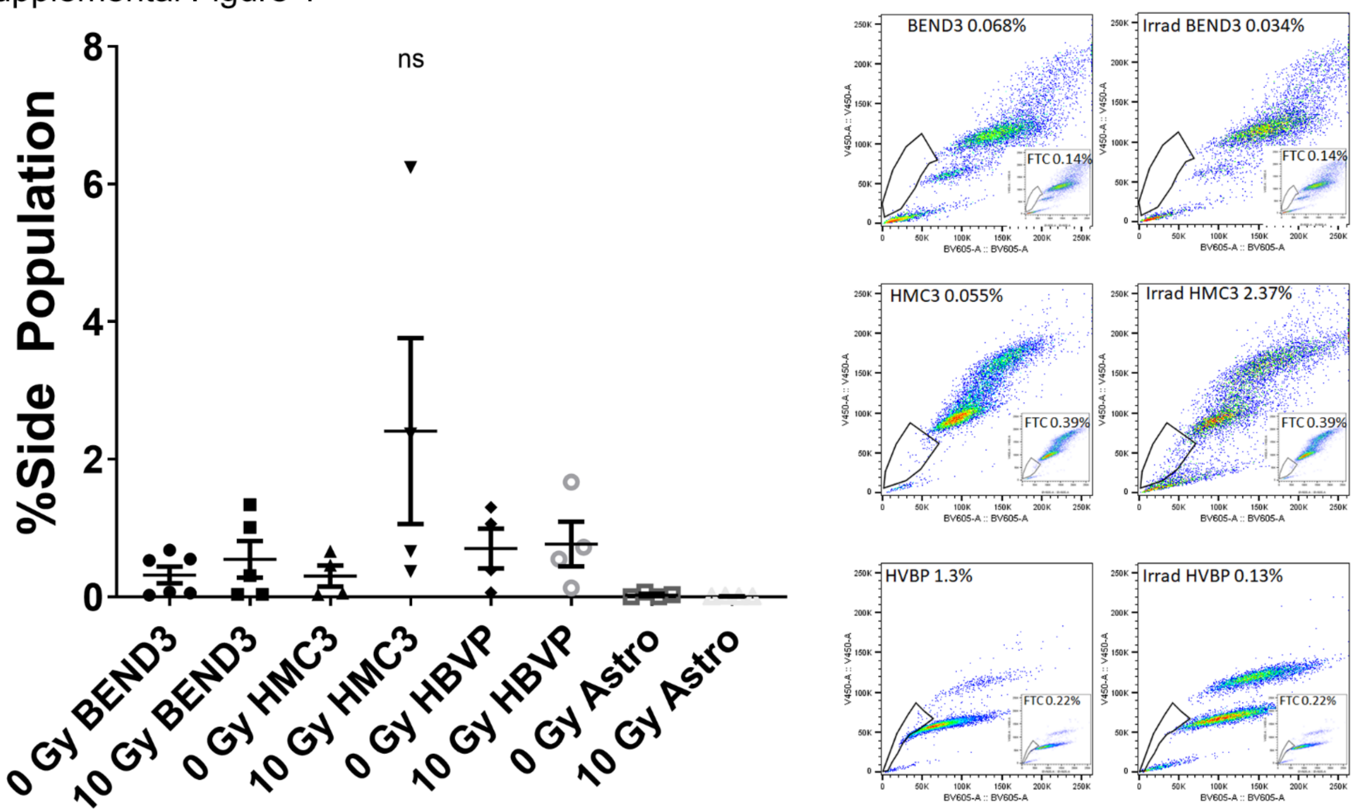


**Supplemental Figure 1. Stromal cells do not have a detectable side population.** Left: Average SP after treatment with either 0 Gy or 10 Gy on the indicated cell type. Error bars indicate SEM. Right: Sample FACS plots show gating for SP.

Supplemental Figure 2

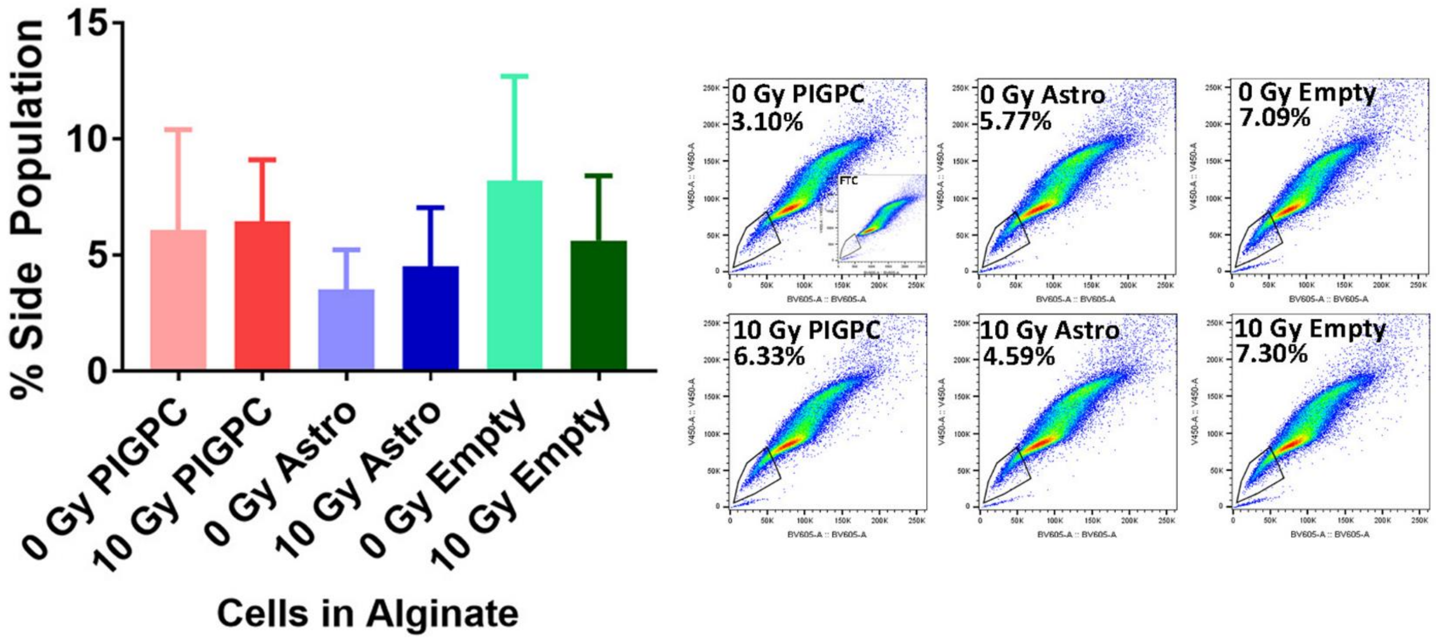


**Supplemental Figure 2. Conditioned media from astrocytes does not stimulate SP of glioma cells.** Left: Average SP of glioma cells after co-culture with sodium alginate beads embedded with either PIGPC, astrocytes (Astro), or no cells (Empty). PIGPC, astrocytes, and empty sodium alginate beads were pretreated with either 0 Gy or 10 Gy. Error bars indicate SEM. Right: Sample FACS plots show gating for SP.

Supplemental Figure 3

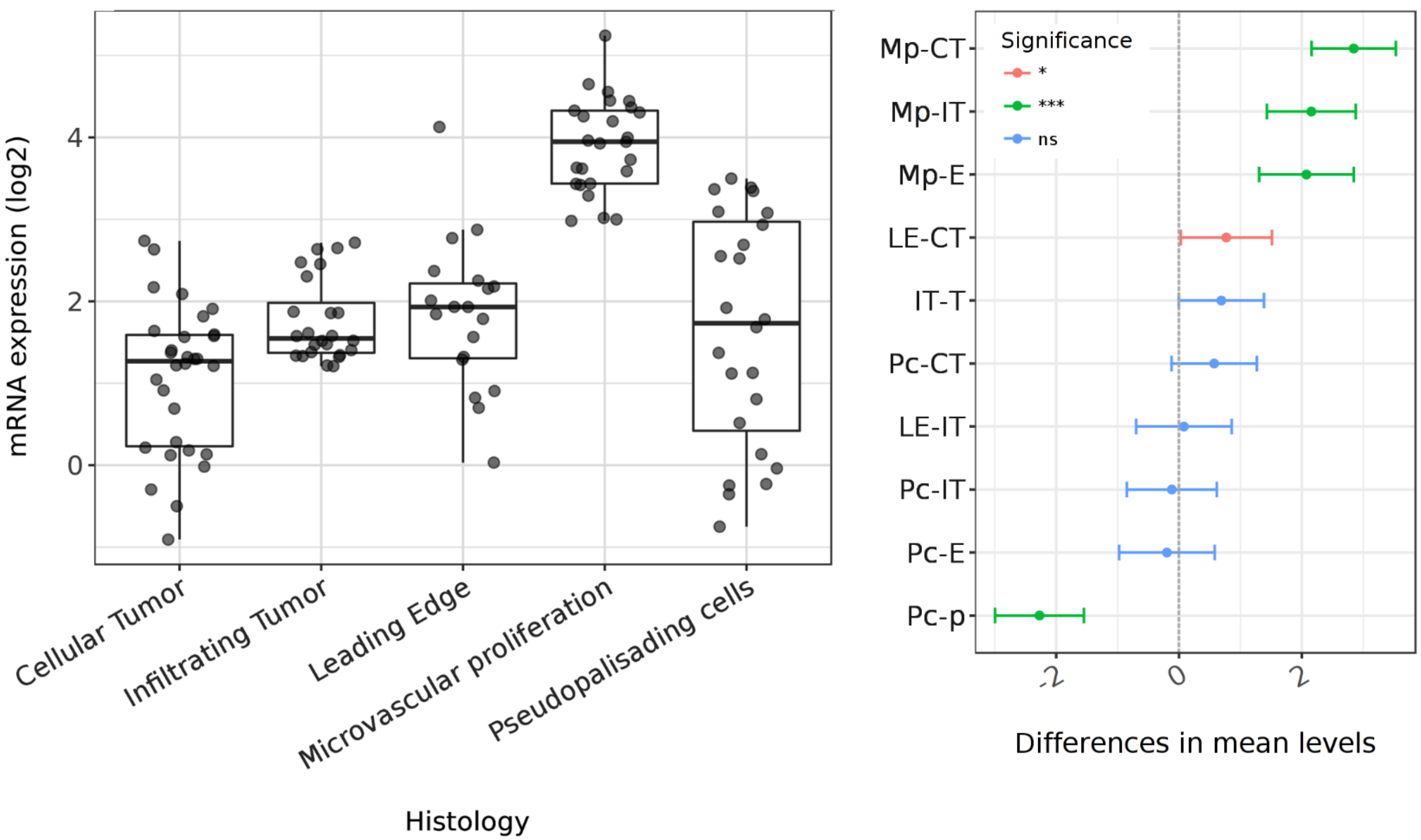


**Supplemental Figure 3. TGM2 is elevated in areas of microvascular proliferation.** TGM2 transcript expression in the indicated histological region of gliomas (left) and statistical analysis (Tukey's Honest Significant Difference, right). Microvascular prolifertion (Mp), Cellular Tumor (CT), Infiltrating Tumor (IT), Leading Edge (LE), Pseudopalisading cells (Pc).

**Supplementary Table 1 Sample Gene Set Enrichment Data of Control versus Irradiated Astrocytes**

| Gene Set Name | NES | FDR q-val |
|---|---|---|
| ZHOU_CELL_CYCLE_GENES_IN_IR_RESPONSE_24HR | 3.3 | 0 |
| WHITEFORD_PEDIATRIC_CANCER_MARKERS | 3.3 | 0 |
| CROONQUIST_IL6_DEPRIVATION_DN | 3.2 | 0 |
| SHEDDEN_LUNG_CANCER_POOR_SURVIVAL_A6 | 3.2 | 0 |
| ZHOU_CELL_CYCLE_GENES_IN_IR_RESPONSE_6HR | 3.1 | 0 |
| GRAHAM_NORMAL_QUIESCENT_VS_NORMAL_DIVIDING_DN | 3.1 | 0 |
| WINNEPENNINCKX_MELANOMA_METASTASIS_UP | 3.1 | 0 |
| KANG_DOXORUBICIN_RESISTANCE_UP | 3.1 | 0 |
| BENPORATH_PROLIFERATION | 3.1 | 0 |
| REACTOME_MITOTIC_M_M_G1_PHASES | 3.0 | 0 |
| REACTOME_DNA_REPLICATION | 3.0 | 0 |
| WHITFIELD_CELL_CYCLE_G2_M | 3.0 | 0 |
| WHITFIELD_CELL_CYCLE_G2 | 3.0 | 0 |
| SARRIO_EPITHELIAL_MESENCHYMAL_TRANSITION_UP | 3.0 | 0 |
| BENPORATH_CYCLING_GENES | 3.0 | 0 |
| REACTOME_CELL_CYCLE | 3.0 | 0 |
| AMUNDSON_GAMMA_RADIATION_RESPONSE | 2.9 | 0 |
| KAUFFMANN_MELANOMA_RELAPSE_UP | 2.9 | 0 |
| TANG_SENESCENCE_TP53_TARGETS_DN | 2.9 | 0 |
| JOHANSSON_GLIOMAGENESIS_BY_PDGFB_UP | 2.1 | 9.43E-05 |
| ZHENG_GLIOBLASTOMA_PLASTICITY_UP | 2.9 | 0 |
| KEGG_MISMATCH_REPAIR | 1.8 | 0.011494 |
| KAUFFMANN_DNA_REPAIR_GENES | 2.2 | 2.74E-05 |
| REACTOME_HOMOLOGOUS_RECOMBINATION_REPAIR_OF_REPLICATION_INDEPENDENT_DOUBLE_STRAND_BREAKS | 2.0 | 6.83E-04 |
| REACTOME_DNA_REPAIR | 1.8 | 0.012718 |
| MACAEVA_PBMC_RESPONSE_TO_IR | -3.3 | 0 |
| WARTERS_RESPONSE_TO_IR_SKIN | -3.1 | 0 |
| SMIRNOV_RESPONSE_TO_IR_6HR_UP | -3.0 | 0 |
| GHANDHI_DIRECT_IRRADIATION_UP | -2.8 | 0 |
| WARTERS_IR_RESPONSE_5GY | -2.8 | 0 |
| PID_P53_DOWNSTREAM_PATHWAY | -2.5 | 0 |
| SCHAVOLT_TARGETS_OF_TP53_AND_TP63 | -2.4 | 0 |

| | | |
|---|---|---|
| AMUNDSON_DNA_DAMAGE_RESPONSE_TP53 | -2.4 | 0 |
| VERHAAK_GLIOBLASTOMA_MESENCHYMAL | -2.3 | 0 |
| SMIRNOV_RESPONSE_TO_IR_2HR_UP | -2.2 | 1.47E-04 |
| ONGUSAHA_TP53_TARGETS | -2.2 | 1.35E-04 |
| KANNAN_TP53_TARGETS_UP | -2.2 | 1.74E-04 |
| RICKMAN_TUMOR_DIFFERENTIATED_WELL_VS_POORLY_DN | -2.1 | 5.26E-04 |
| INGA_TP53_TARGETS | -2.1 | 8.94E-04 |

**Supplementary Table 2: Changes in Matrix Proteins From Astroctyes After 10 Gy**

| Protein | Significance | Percent Coverage | Peptides | Unique Peptides | Gene Name | Ratio to Control | Accession |
|---|---|---|---|---|---|---|---|
| Pentraxin-related protein PTX3 | 85.43 | 13 | 3 | 3 | PTX3 | 4.53 | P26022 |
| Transglutaminase 2/Protein-glutamine gamma-glutamyltransferase 2 | 138.9 | 47 | 29 | 29 | TGM2 | 4.16 | P21980 |
| Nestin | 57.51 | 9 | 11 | 11 | NES | 3.24 | P48681 |
| Fibulin-1 | 117.03 | 42 | 19 | 19 | FBLN1 | 2.94 | P23142 |
| Nicastrin | 59.86 | 4 | 3 | 3 | NCSTN | 2.04 | Q92542 |
| Matrilin-2 | 50.85 | 2 | 2 | 2 | MATN2 | 2.02 | O00339 |
| Matrix-remodeling-associated protein 7 | 50.23 | 12 | 2 | 2 | MXRA7 | 1.82 | P84157 |
| Endonuclease domain containing 1 protein | 63.61 | 21 | 6 | 6 | ENDOD1 | 1.71 | O94919 |
| Glypican-4 | 71.37 | 6 | 2 | 2 | GPC4 | 1.69 | O75487 |
| ADAMTS-like protein 4 | 79.86 | 3 | 3 | 3 | ADAMTSL4 | 1.61 | Q6UY14 |
| Prolyl endopeptidase FAP | 51.57 | 21 | 14 | 14 | FAP | 1.57 | Q12884 |
| Fibronectin | 110.19 | 66 | 163 | 163 | FN1 | 1.24 | P02751 |
| Laminin subunit beta 2 | 56.62 | 4 | 5 | 5 | LAMB2 | 1.18 | P55268 |
| Collagen alpha-1(IV) chain | 65.43 | 3 | 6 | 4 | COL4A1 | 1.04 | P02462 |
| Collagen alpha-1(XI) chain | 69.11 | 5 | 6 | 6 | COL11A1 | 0.55 | P12107 |
| Cathepsin B | 68.45 | 39 | 10 | 10 | CTSB | 0.52 | P07858 |
| Lactadherin | 57.4 | 22 | 6 | 6 | MFGE8 | 0.51 | Q08431 |
| Beta-2-microglobulin | 60.15 | 18 | 2 | 2 | B2M | 0.5 | P61769 |
| SPARC | 71.04 | 23 | 5 | 5 | SPARC | 0.44 | P09486 |
| Microfibrillar-associated protein 2 | 54.91 | 10 | 2 | 2 | MFAP2 | 0.42 | p55001 |
| Collagen alpha-3(VI) chain | 76.17 | 28 | 64 | 64 | COL6A3 | 0.38 | P12111 |
| Fibrillin-2 | 61.46 | 5 | 12 | 9 | FBN2 | 0.37 | P35556 |
| EGF-like repeat and discoidin I-like domain-containing protein 3 | 61.14 | 13 | 5 | 5 | Edil3 | 0.29 | O43854 |
| Collagen alpha-1(I) chain | 87.25 | 7 | 6 | 5 | COL1A1 | 0.24 | P02452 |
| Biglycan/PGS1 | 62.63 | 12 | 3 | 3 | BGN | 0.16 | P21810 |
| High mobility group protein B1 | 76.97 | 15 | 4 | 2 | HMGB1 | 0.08 | P09429 |
